# Synapse-specific direction selectivity in retinal bipolar cell axon terminals

**DOI:** 10.1101/2020.10.12.335810

**Authors:** Akihiro Matsumoto, Weaam Agbariah, Stella Solveig Nolte, Rawan Andrawos, Hadara Levi, Shai Sabbah, Keisuke Yonehara

## Abstract

The ability to encode the direction of image motion is fundamental to our sense of vision. Direction selectivity along the four cardinal directions is thought to originate in direction-selective ganglion cells (DSGCs), due to directionally-tuned GABAergic suppression by starburst cells. Here, by utilizing two-photon glutamate imaging to measure synaptic release, we reveal that direction selectivity along all four directions arises earlier than expected, at bipolar cell outputs. Thus, DSGCs receive directionally-aligned glutamatergic inputs from bipolar cell boutons. We further show that this bouton-specific tuning relies on cholinergic excitation and GABAergic inhibition from starburst cells. In this way, starburst cells are able to refine directional tuning in the excitatory visual pathway by modulating the activity of DSGC dendrites and their axonal inputs using two different neurotransmitters.

## Results

Retinal bipolar cells are excitatory glutamatergic interneurons that transfer visual information from photoreceptors to the inner retina, where ganglion cells form the main output to the brain (*1*). There are 14 molecularly distinct types of bipolar cell in the mouse retina, each carrying different types of visual information, including luminance change, temporal kinetics, and chromatic sensitivity (*1–3*). Information about the direction of moving stimuli is thought to be encoded by a specialized class of retinal ganglion cells, DSGCs, which receive inputs from GABAergic/cholinergic starburst amacrine cells as well as glutamatergic bipolar cells. DSGCs respond to one of the four cardinal directions (dorsal, ventral, nasal, and temporal) with maximal spiking in their preferred direction, but only minimal spiking in the opposite (null) direction (*4*). Thus, DSGCs embody the retinal coordinates of motion computation (*5*). A body of literature supports the consensus that cardinal direction selectivity emerges first at DSGC dendrites, due to directionally-tuned GABAergic inhibition from starburst cells (*4, 6–8*). Indeed, studies that have imaged extracellular glutamate at DSGC dendrites without identifying the types of input bipolar cells (*9, 10*), or intracellular calcium in axon terminals of a small range of specific bipolar cell types (*9, 11*), have failed to identify directionally-tuned activity in bipolar cells. However, there is anecdotal evidence to the contrary (*12*), and none of the reports have incorporated a comprehensive analyses of functionally classified bipolar cell types.

Motivated by two recent findings that 1) DSGC dendrites receive locally tuned cholinergic inputs from starburst cells (*13*) and 2) alpha-7 nicotinic acetylcholine receptors (α7-nAChRs) are selectively expressed in type 7 (T7) ON and type 2 (T2) OFF bipolar cell (*14*), we tested the potential role of cholinergic signaling in the motion-related modulation of these specific bipolar cell types outputs. Subretinal injection of an adeno-associated virus (AAV) vector of serotype 8BP/2 (*15*), containing a CAG promoter, preferentially labeled T7, T2, and rod bipolar cells (Xin Duan, personal communication on CAG promoter tropism; Fig. 1A). This was confirmed by the depths to which labeled bipolar cell axon terminals projected in mice in which starburst cell processes were labeled with tdTomato (Fig. 1, B and C; fig. S1) (*1, 3, 14, 16*). This strategy allowed us to target these bipolar cell types with a glutamate sensor SF-iGluSnFR.A184S for imaging light-evoked glutamate release from axon terminals of T7 and T2 bipolar cells (Fig. 1, D and E; fig. S2, A to D). A number of neighboring axonal boutons exhibited correlated noise during a static flash (Fig. 1D; fig. S2, I to K), indicating that they belonged to the same bipolar cell (*2, 17*). Strikingly, motion stimuli revealed directional tuning in a fraction of boutons (Fig. 1, E to G), which displayed heterogenous direction preference (Fig. 1H; direction selective index (DSI), 0.29 ± 0.19, 1108 ON boutons; fig. S2E). There was no bias towards a specific direction (Fig. 1, I and J; fig. S2, F and G; *p* > 0.1, Hodges-Ajne test), but instead clusters along the four cardinal directions associated with ON-OFF DSGC firing patterns (Fig. 1J; *p* = 0.037, number of modes = 4, Silverman’s test; fig. S2H) (*4, 5*).

**Fig. 1.**
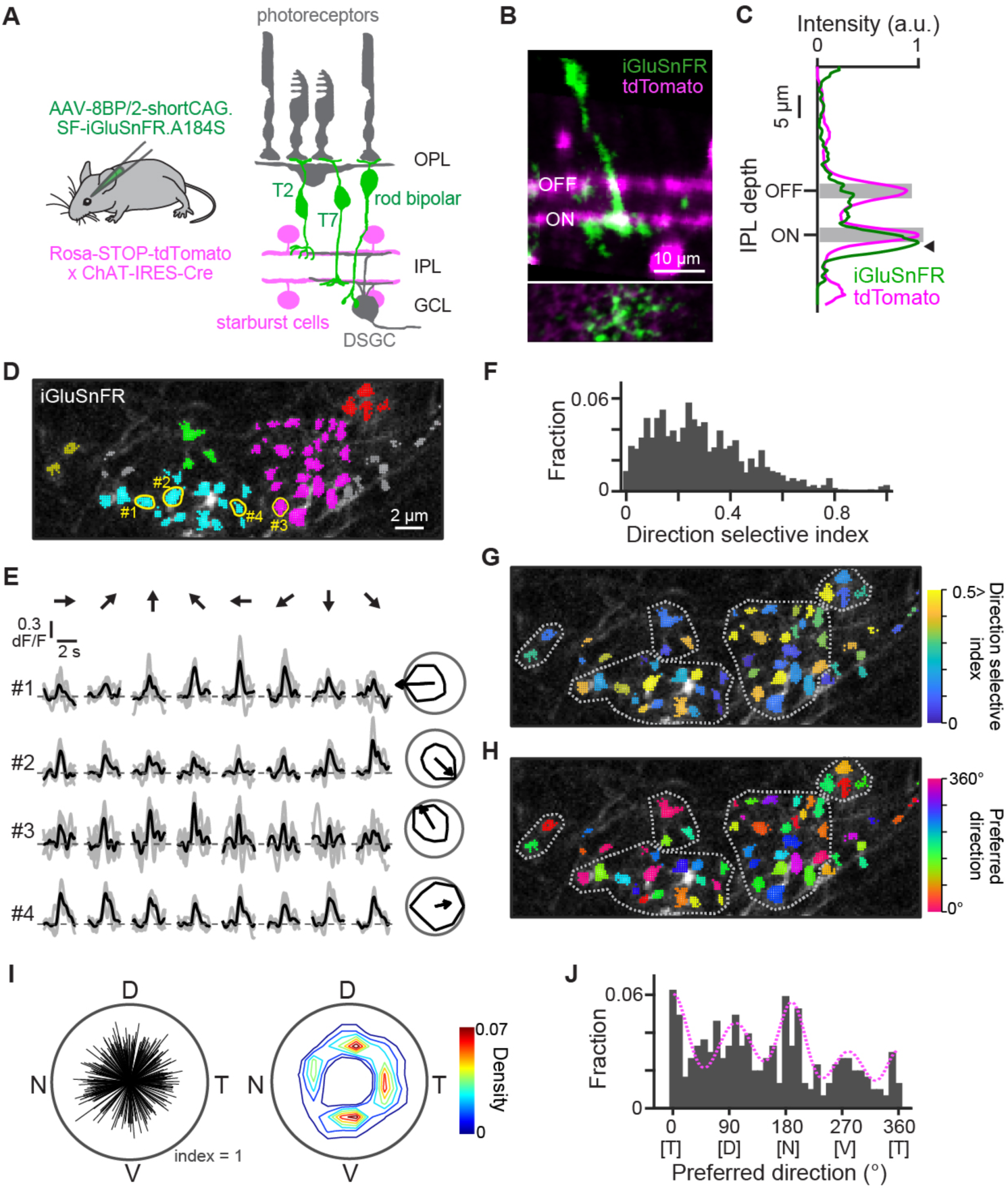
Type 7 bipolar cell axon terminals display cardinal direction selectivity. (**A**) Glutamate sensor SF-iGluSnFR.A184S was selectively targeted to type 2 (T2), type 7 (T7), and rod bipolar cells (green) by subretinal injection of AAV-8BP/2 with a CAG promotor into Rosa-STOP-tdTomato x ChAT-IRES-Cre mice. Magenta, starburst cells labeled by tdTomato. (**B**) Example cross-sectional view (top) and tangential view (bottom; at the depth of the arrowhead in C) of two-photon images of an SF-iGluSnFR.A184S-labeled T7 bipolar cell (green) and tdTomato-labeled starburst cells (magenta). (**C**) The cross-sectional intensity of iGluSnFR (green) and tdTomato (magenta) fluorescence in B. Arrowhead, depth in the inner plexiform layer (IPL) at which two-photon glutamate imaging was performed. Gray bands, depth of ON and OFF tdTomato signals. (**D**) Example field-of-view during glutamate imaging. Axonal boutons with high noise correlation are indicated the same color (Materials and Methods; fig S2, I to K). Gray, unclassified boutons. Yellow lines, example regions of interest for E. (**E**) Glutamate signals during motion stimuli (gray, individual trials; black, average) and directional tuning (right) in tuned (#1-#3) and untuned (#4) boutons shown in D. (**F**) DSI histograms of ON (n = 1108) boutons. (**G and H**) DSI (G) and preferred direction (H) of individual boutons. Dotted gray lines, rough border of each cell. (**I**) Left, polar plot of directional tuning in individual ON boutons with >0.3 DSI (305 boutons; length, DSI; angle, preferred direction). Right, density plot of the left panel. (**J**) Histogram of the preferred direction of ON boutons. Magenta curve, fitted Gaussian function.

Axon terminals of bipolar cells are modulated by different types of amacrine cells (*2, 18, 19*), which might be responsible for the direction-selective glutamate release that we observed from bipolar cell terminals. Given the specific expression and conductance of α7-nAChRs in T7 and T2 bipolar cells (*14*), one potential mechanism is enhancement of glutamate release in response to the preferred direction by cholinergic starburst cells (*20*). Alternatively, GABAergic inhibition from wide-field amacrine cells (*21*), driven by voltage-gated sodium channels (Na_V_), could give rise to suppression in response to the null direction motion. To test these possibilities, we mapped the effects of pharmacological manipulation on individual boutons along the axon terminals of single T7 bipolar cells (Fig. 2, A and B). We found that blockade of both α7-nAChR by α-Bungarotoxin (α-Bgtx) and Na_V_ by tetrodotoxin (TTX) diminished the tuning of direction-selective bipolar cell boutons.

**Fig. 2.**
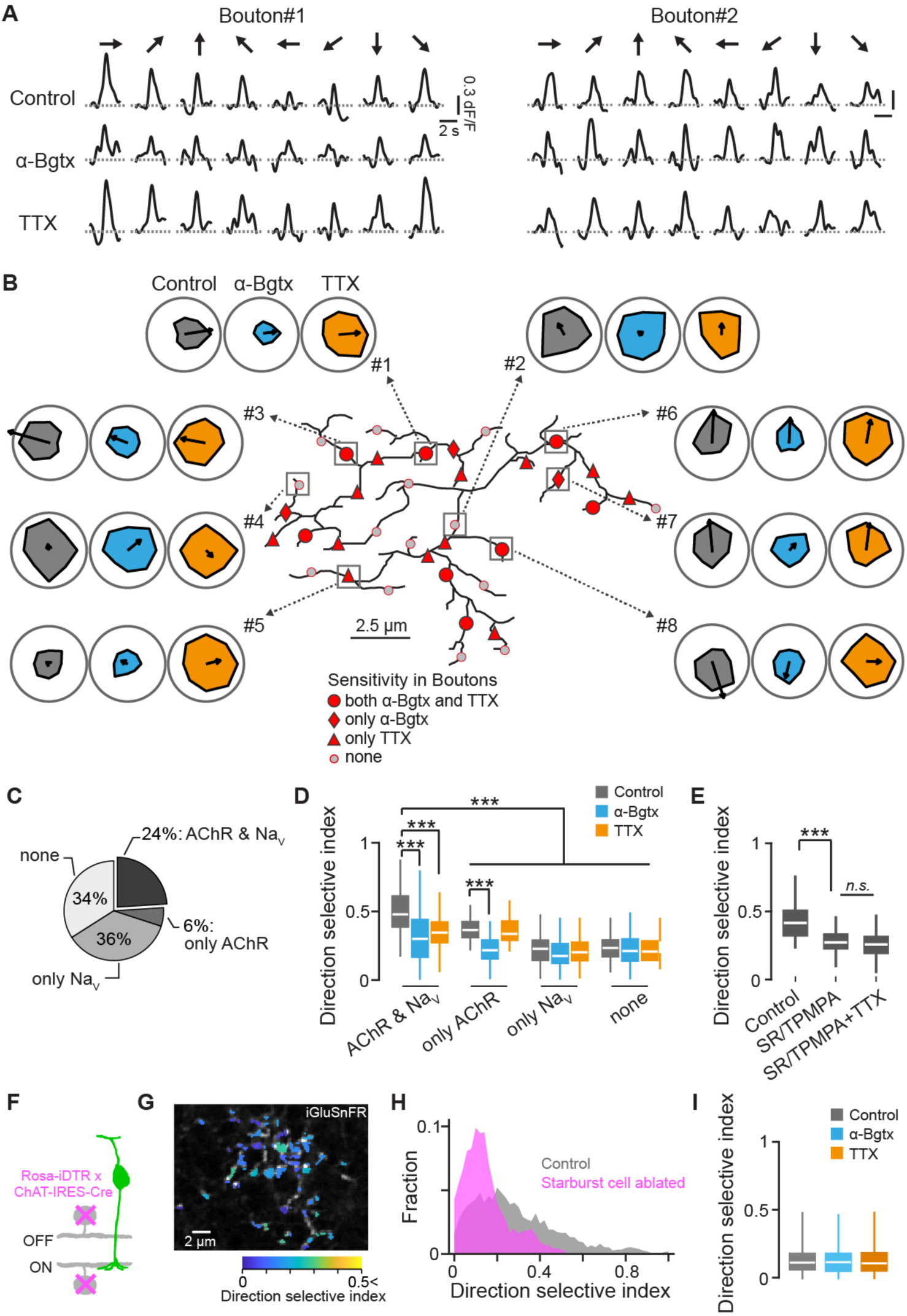
Glutamate release from T7 bipolar cells is regulated at individual axonal boutons. (**A**) Example average responses of T7 bipolar cell axonal boutons to motion stimuli before and during pharmacological blockade of α7-nAChR by α-Bgtx and Na_V_ by TTX (three trials). The location of boutons #1 and #2 are indicated in B. (**B**) Directional tuning of individual boutons on axonal branches (black lines) of a T7 bipolar cell. The effects of blockade are indicated by polar plots (gray, control; cyan, α-Bgtx; orange, TTX). Different sensitivities of boutons are depicted by red circles (sensitive to both α-Bgtx and TTX), red diamonds (sensitive only to α-Bgtx), red triangles (sensitive only to TTX), and gray circles (insensitive). (**C**) Populations of bouton types with distinct sensitivity determined from five T7 bipolar cells. Sensitivity was calculated by using responses to a static flash stimulus (fig S3, A to C). (**D**) Summary of DSI changes due to blockade in the four bouton types (128 both, 56 α-Bgtx-only, 181 TTX-only, 194 neither). (**E**) DSI changes in 58 boutons due to GABA receptor blockade (SR95531 + TPMPA) and additional application of TTX. (**F**) Starburst cells (magenta) in Rosa-iDTR x ChAT-IRES-Cre mice were selectively ablated by intravitreal injection of diphtheria toxin. (**G**) DSI for labeled T7 bipolar cell axons in starburst cell-ablated retinas. (**H**) DSI histograms in control (gray; 493 boutons, 4 retinas) and starburst cell-ablated retinas (magenta; 386 boutons, 5 retinas). (**I**) Comparisons of DSI in control (gray), α-Bgtx (cyan), and TTX (orange) conditions in starburst cell-ablated retinas (no significance). ***, *p* < 0.001. Mann-Whitney-Wilcoxon test for Fig. 2D; Wilcoxon signed-rank test for Fig. 2E.

These experiments revealed that boutons could be classified into four groups based on their sensitivity to pharmacological manipulation (Fig. 2C; fig. S3, A to D): 24% were sensitive to both α-Bgtx and TTX; 6% and 36% were sensitive only to α-Bgtx or TTX, respectively; and 34% of were sensitive to neither. Boutons that were sensitive to both α7-nAChR and Na_V_ blockade showed significantly higher directional tuning under control conditions (mean DSI, 0.29 ± 0.19; Fig. 2D), compared to other boutons. In TTX-sensitive boutons, the effect of Na_V_ blockade was occluded by pre-application of SR95531 and TPMPA, which block GABA_A_ and GABA_C_ receptors, respectively (Fig. 2E). This reveals that GABAergic inhibition by Na_V_-expressing wide-field cells, rather than Na_V_-expressing bipolar cell types (*18, 22, 23*), mediates directional tuning.

Directional tuning in α7-nAChR-blocked boutons was further reduced by subsequent application of SR95531 and TPMPA (fig. S3E) or TTX (fig. S3F), suggesting that GABAergic inhibition and cholinergic excitation are mechanistically independent. Furthermore, glutamate release from axonal boutons belonging to the same bipolar cell was highly correlated during static stimuli, but decorrelated during moving stimuli (fig. S3G). This motion-induced decorrelation was occluded by α-Bgtx and TTX, suggesting that glutamate release from T7 and T2 bipolar cells can be driven by dendritic inputs from photoreceptors, and modulated by the cholinergic excitation and GABAergic inhibition (fig. S3, G and H).

To examine the causal contribution of starburst cells to direction-selective glutamate release from bipolar cells, we selectively ablated starburst cells by intravitreal injection of diphtheria toxin to Rosa-iDTR × ChAT-IRES-Cre mice (Fig. 2F; fig S4, A to F). Direction selectivity in both T7 and T2 bipolar cells reduced significantly (Fig. 2, G and H, *p* = 6.82×10^−37^; fig S4, J to L). Blockade of α7-nAChRs did not affect direction selectivity in these ablated retinas, confirming that starburst cells mediate the cholinergic-dependent component of direction selectivity in bipolar cell terminals (Fig. 2I, cyan; fig S4, G and H). We further found that starburst cell ablation also occluded the effect of TTX on direction selectivity (Fig. 2I, orange; fig S4, H and I), suggesting that the contribution of wide-field cells to directional tuning is dependent on starburst cells.

To anatomically confirm the involvement of starburst and wide-field amacrine cells in bipolar cell computation of motion stimuli, we used a serial block-face electron microscopy (SBEM) dataset (*24*). We first reconstructed all ON and OFF bipolar cells that formed synapses with ON-OFF DSGC cells (fig. S5, A to D) (*13, 24, 25*), then focused on one of the reconstructed T7 bipolar cells (fig. S5E) for further tracing of synapses from amacrine cells onto this T7 bipolar cell. We identified a total of 112 amacrine cell-to-bipolar cell synapses and reconstructed the amacrine cell dendritic processes (Fig. 3, A to C). These reconstructed amacrine cells fell into two categories: wide-field cells (31%) with long axon-like processes showing little, if any, branching (Fig. 3B and D), and non-wide-field cells (69%) with highly-branched processes (Fig. 3, C and D; fig. S5, H to J). None of the non-wide-field cells were identified as starburst cells, which is consistent with previous reports (*11, 26*) and suggests that cholinergic signaling from starburst cells to T7 bipolar cells might be mediated by recently-identified non-synaptic, locally-tuned forms of transmission (*13*) rather than by clear synaptic structures. Of the terminal branches with output ribbon synapses that we observed, 20% received synapse from wide-field cells, 20% from both wide-field and non-wide-field cells, and 12% from only non-wide-field cells (Fig. 3E). 48% branches did not receive any inputs.

**Fig. 3.**
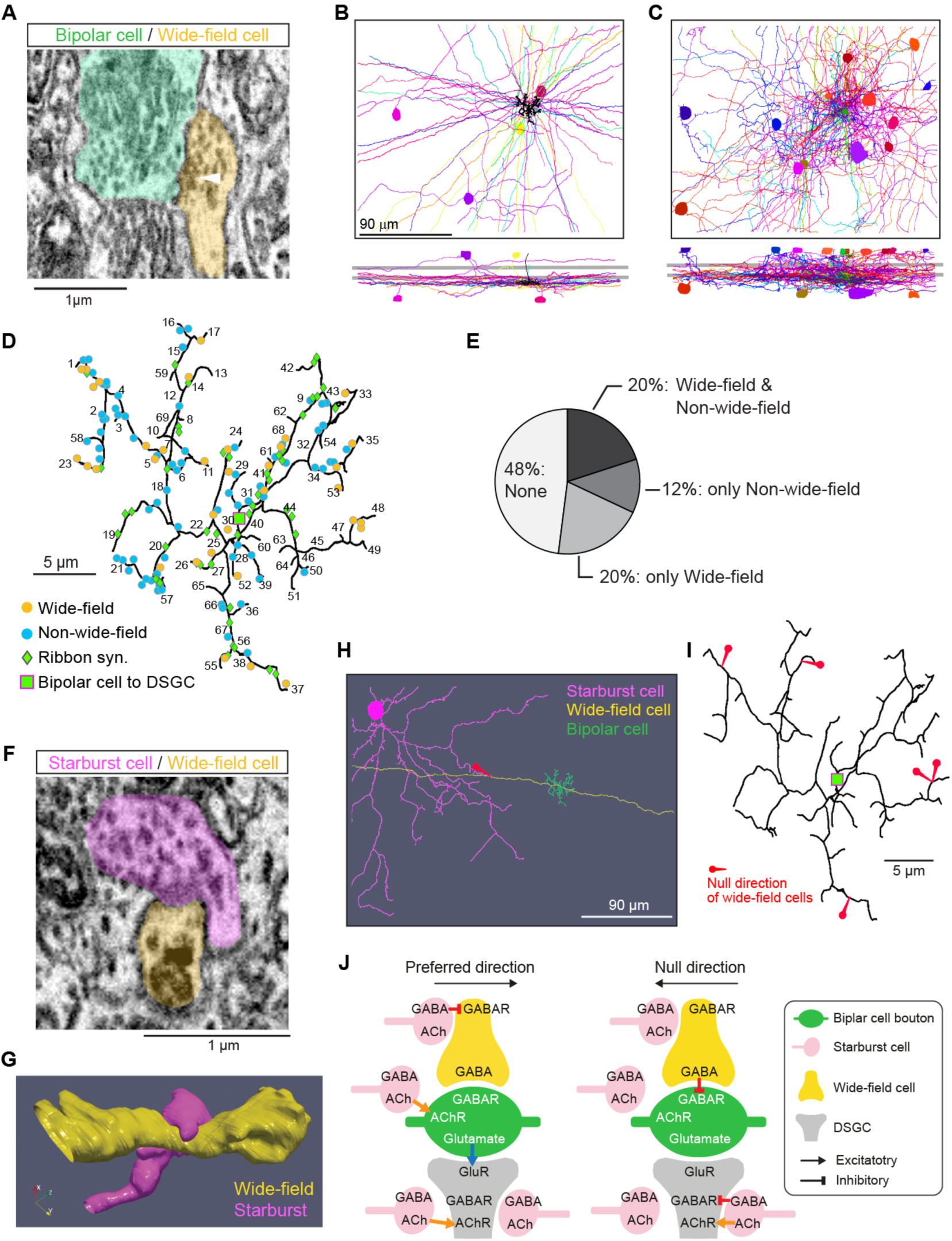
Amacrine cells form synapses with axon terminal branches of a T7 bipolar cell (**A**) An example synapse between a wide-field amacrine cell (yellow) on a terminal branch of a T7 bipolar cell (light green). Arrowhead indicates a presynaptic site of the amacrine cell. (**B and C**) *en face* (top) and orthogonal (bottom) views of partially reconstructed wide-field (B) and non-wide-field (C) amacrine cells that are presynaptic to the terminal branches of a T7 bipolar cell. The bipolar cell axon terminal is shown in black close to the volume center. (**D**) Axon terminal branches of a T7 bipolar cell (black lines), indicating the numerical IDs of each branch. Circles, location of the input synapses made by wide-field (yellow) and non-wide-field (cyan) amacrine cells. Green diamonds, output dyad ribbon synapses from bipolar cell. Green square, a dyad ribbon synapse from the bipolar cell onto its apposing DSGC. (**E**) The population of input types on the T7 bipolar cell terminal branches that have ribbon synapses in D (52% of the branches). 48% branches did not receive any inputs (“None”). (**F and G**) Electron micrograph (F) and 3D reconstruction (G) of a starburst-to-wide-field cell synapse (magenta, starburst cell; yellow, wide-field cell). (**H**) The wiring between a starburst cell (magenta), wide-field cell (yellow), and T7 bipolar cell (green). The vector connecting the starburst cell soma to the starburst-to-wide-field cell synapse (red arrow) indicates an estimation of the wide-field cell’s null direction. (**I**) Mapping of estimated null directions of wide-field cells (red arrows) on the T7 bipolar cell axon terminal (black). (**J**) Schematic of potential circuit mechanisms for direction-selective glutamate release. In the preferred direction (left), the bipolar cell bouton (green) and the DSGC dendrite (gray) receives excitation mediated by ACh (orange arrow) nonsynaptically transmitted from a starburst cell (middle and bottom red), and the wide-field cell (yellow) is inhibited by GABA released from a starburst cell (top red). In the null direction (right), a wide-field cell inhibits the bipolar cell bouton, and a DSGC dendrite is inhibited and excited by GABA and ACh, respectively, co-released from a starburst cell (bottom red) as previously demonstrated (*7*).

We subsequently searched for connections between wide-field cells and starburst cells by scanning the axons of wide-field cells that formed synapses with the reference T7 bipolar cell axon terminals. We found synapses from starburst cells to the axons of wide-field cells, but not in the vicinity of the ribbon synapse between the bipolar cell and DSGC (Fig. 3, F to H). Interestingly, the estimated preferred directions of wide-field cells varied among the terminal branches of the T7 bipolar cell arbor (Fig. 3I), indicating that GABAergic inhibition of T7 cell boutons is tuned to different directions (Fig. 1I).

Together, these anatomical, genetic, and pharmacological analyses suggest two potential circuit mechanisms that could establish direction-selective glutamate release by T7 bipolar cells (Fig. 3J). The first is preferred direction enhancement of release via α7-nAChR activation at the bipolar cell axon terminal (Fig. 3J, left). The second is null direction inhibition of release via GABAergic Na_V-_expressing wide-field cells (Fig. 3J, right). In the null direction, the lack of ACh input to the bipolar cell terminal and the induction of GABAergic inhibition from wide-field cells would result in suppressed glutamate release.

We next sought to test whether the directional tuning in T7 and T2 bipolar cell terminal boutons is received by ON-OFF DSGCs by targeting DSGCs labeled in the retinas of Cart-IRES-Cre mice (*27*) with Cre-dependent AAVs expressing iGluSnFR. Strikingly, two-photon imaging of iGluSnFR-labeled DSGC dendrites (Fig. 4, A and B; fig. S6, A to F) revealed that DSGCs receive both directionally-tuned and untuned glutamate inputs (Fig. 4, B to D). To examine T7 and T2 bipolar cell inputs, we initially performed unsupervised clustering to identify response kinetics (*2, 18, 28, 29*) (fig. S6, G to K). We then identified six ON groups along the ON arbor (Fig. 4E, F; G1-G6) and six OFF groups along the OFF arbor (fig. S6, I to K; G7-G12) of DSGC dendrites. Of these, groups G3 (ON) and G11 (OFF) were selectively modulated by α7-nAChR block (Fig. 4F; fig. S6L), suggesting that they correspond to inputs from T7 and T2 bipolar cells, respectively.

**Fig. 4.**
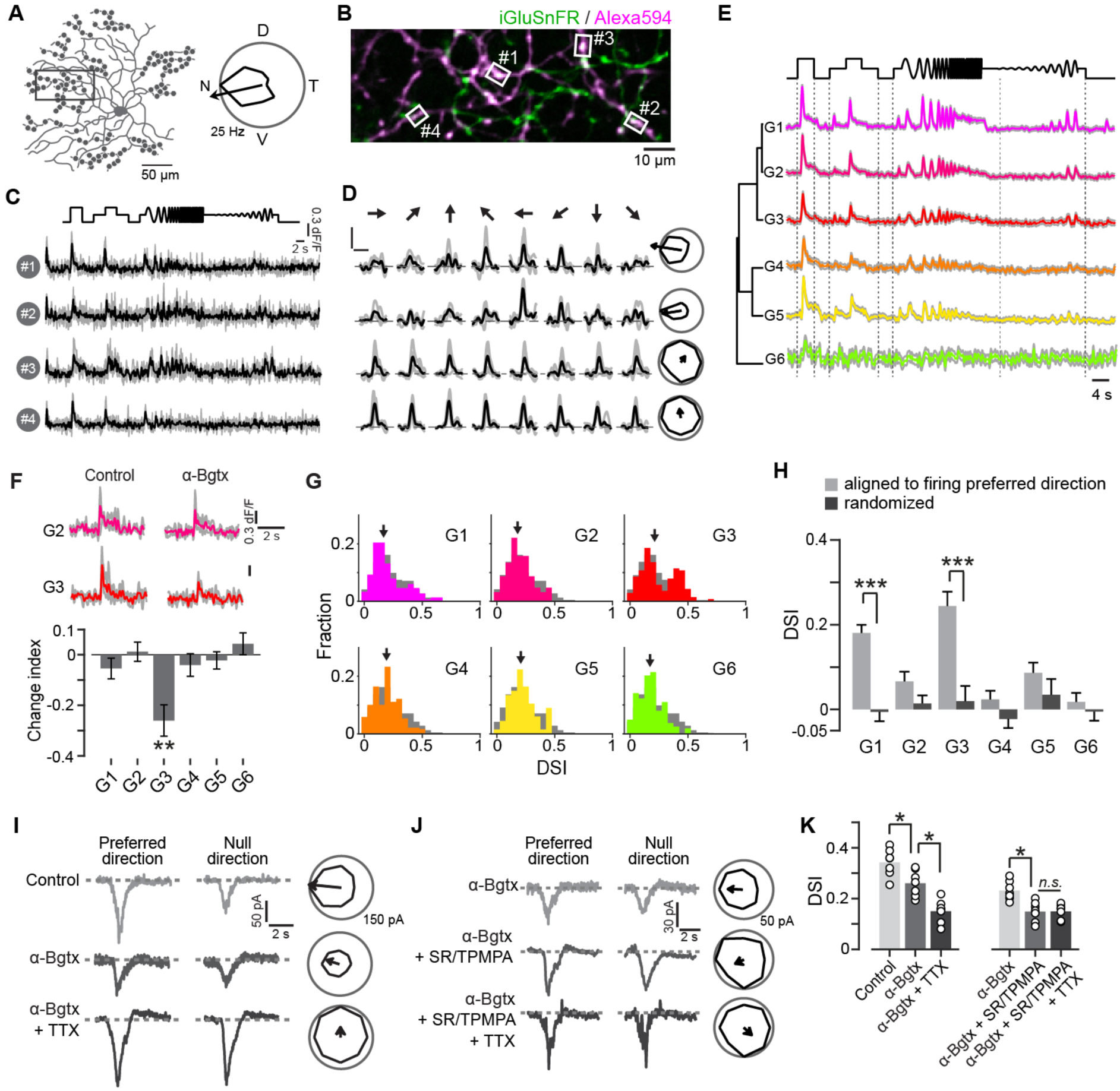
DSGCs receive direction preference-matched glutamatergic inputs from bipolar cells (**A**) The reconstructed morphology of an example ON-OFF DSGC (left) that was filled with Alexa 594 for two-photon imaging. Gray dots depict sites of glutamatergic input. Directional tuning of firing in this neuron (right). Arrow, vector sum indicating preferred direction. T, temporal. D, dorsal. N, nasal. V, ventral. (**B**) An example two-photon image of the field of view indicated in the gray rectangle in A. (**C and D**) Glutamate signals during modulating spot (C) and motion stimuli (D) in regions of interests (white squares in B). Gray and black, each trial and average, respectively. (**E**) Average modulating spot-evoked responses in the six identified ON groups (G1-6). Gray shade, SD. (**F**) Top, example glutamate signals from G2 (top) and G3 (bottom) groups in response to static flash (2 s duration; 100% contrast) before (left) and after the blockade of α7-nAChR by α-Bgtx. Gray and colored, each trial and average, respectively. Bottom, change index showing the effects of α-Bgtx application on response amplitudes. 46 G1, 38 G2, 26 G3, 41 G4, 45 G5, 27 G6 inputs. (**G**) DSI histogram of all inputs (gray) and 6 ON groups (colored). Arrows, median. (**H**) Comparison between DSI determined along the preferred direction of the recipient DSGC (light gray) and DSI determined along a randomly selected direction (dark gray). (**I**) EPSCs recorded from an example ONOFF DSGC evoked by the leading edge of a spot moving in the preferred and null direction, and their directional tuning in control conditions and following α-Bgtx and TTX application. (**J**) EPSCs and their directional tuning following additive blockade by α-Bgtx, SR95531/TPMPA, and TTX. (**K**) Summary of DSI changes due to pharmacological manipulations. Bars and circles, averages and 7 individual cells, respectively. ***, *p* < 0.001; **, *p* < 0.01; *, *p* < 0.05. Wilcoxon signed-rank test.

Among the twelve groups, distributions of DSI were skewed toward higher values in groups G3 and G11 (Fig. 4G; skewness and median: 0.42, 0.21 in G3; 0.44, 0.26 in G11; 0.12, 0.13 in all inputs; fig. S6, M to P), indicating that they include subsets of inputs with higher directional tuning. These directional tunings were diminished by α7-nAChR and Na_V_ blockade (fig. S7A). In particular, the G3 group displayed a heterogeneous response to these blockers, similar to that of T7 bipolar cells (fig. S7, B and C): 21% of inputs were sensitive to both TTX and α-Bgtx (compared to 24% of T7 bipolar axonal boutons; Fig. 2C), 7% were sensitive only to α-Bgtx, 28% only to TTX, and 44% were insensitive to both. Intriguingly, the DSI along the preferred direction for each DSGC was significantly higher in G1 and G3, compared to the low DSI along a randomized direction (Fig. 4H; fig. S7, D to F). When considered alongside the results from axon terminal imaging, these data suggest that DSGCs selectively receive glutamatergic inputs matched to their preferred direction from among a heterogeneously-tuned range of T7 and T2 bipolar cell boutons.

Finally, we tested the contribution of T7 and T2 bipolar cell inputs to the tuning of DSGCs using two-photon targeted patch-clamp recordings from EYFP-labeled DSGCs in retinas isolated from Oxtr-T2A-Cre; Thy1-STOP-EFYP mice (*13*) (Fig. 4, I and K; fig. S7, G and H). Blockade of Na_V_ and α7-nAChR reduced the direction-selectivity of excitatory postsynaptic currents (EPSCs) recorded in DSGCs. TTX-sensitive tuning reduction was occluded by pre-application of GABA receptor blockers (Fig. 4, J and K), revealing that this inhibitory effect was mediated by Na_V-_dependent wide-field cells. The amplitude of inhibitory inputs to DSGCs in the null direction was also decreased by α-Bgtx (fig. S7, I and J), suggesting that the dendritic processes of starburst cells are excited by T7 and T2 bipolar cells, as previously shown (*30, 31*).

## Discussion

Previous work has established the view that starburst cell-to-ganglion cell synapses are the only sites involved in computation of cardinal motion direction selectivity in the retina (*9–11*). Our results show instead that cardinal direction selectivity first emerges in the axon terminals of bipolar cells, and that this axonal direction selectivity is due to cholinergic excitation from starburst amacrine cells and GABAergic null direction suppression from wide-field amacrine cells. These fine microcircuit mechanisms are implemented at individual synapses in such a precise way that directionally-tuned outputs can be transmitted to functionally aligned DSGC types. Future studies could investigate the developmental mechanisms that allow starburst cells to align the tuning of both input axon terminals and recipient dendrites.

Although synapses from T7 and T2 bipolar cells to DSGCs are not abundant (3.4% and 15.3%, respectively), our findings suggest that they contribute to direction selectivity in ganglion cells using at least two distinct mechanisms. First, the tuning of these bipolar cell boutons shapes directionally-sensitive excitatory inputs to DSGCs (Fig. 4, I to K). Second, these bipolar cells drive GABA releases from starburst cells (fig. S7, I and J) (*30, 31*). This concept is supported by a previous study showing that genetic perturbation of T7 bipolar cells weakens the tuning of ONOFF DSGCs (*32*).

Intriguingly, previous DSGC glutamate imaging studies and bipolar cell axon terminal calcium imaging studies have failed to identify directionally-tuned bipolar terminals. This discrepancy could be due to imaging populations versus specific cell types, use of larger regions of interest that blend different tuning types (*9, 10*) compared to those in this study (fig. S6F), or underestimates of the tuning strength due to potential nonlinear Ca^2+^ signals in the axon terminals (*33*) in a previous study (*11*). Importantly, the response amplitudes in our glutamate imaging experiments were set within a linear range to faithfully capture release tuning (fig. S2M; *R*^2^, 0.98).

What is the advantage of encoding all of four cardinal directions, rather than only one, in individual T7 and T2 bipolar cells? Because visual space is represented by a mosaic of bipolar cells of the same type (*1*), multiplexed axonal direction selectivity in T7 and T2 bipolar cells would be an efficient way of encoding all four directions for both ON and OFF responses at every point in the visual space without having extra cell types.

## Acknowledgments

We thank David Berson for providing us with an SBEM volume with traced DSGCs and bipolar cells, Xin Duan for sharing unpublished findings on viral tropism, Zoltan Raics for developing our visual stimulation system, and Bjarke Thomsen and Misugi Yonehara for technical assistance. We also thank Antonia Drinnenberg, Karl Farrow, and Stuart Trenholm for critical comments on an early version, and Lesley Anson for comments on a later version, of the manuscript.

## Funding

VELUX FONDEN Postdoctoral Ophthalmology Research Fellowship (27786) to A.M., Lundbeck Foundation (DANDRITE-R248-2016-2518), Novo Nordisk Foundation (NNF15OC0017252), Carlsberg Foundation (CF17-0085), and European Research Council Starting (638730) grants to K.Y.

## Author contributions

A.M. and K.Y. conceived and designed the experiments and analyses. A.M. performed all physiological experiments. A.M. and S.S.N. performed immunohistochemistry and confocal scanning. K.Y. designed viral vectors and performed eye injections. A.M. analyzed the physiological and confocal data. W.A., R.A., H.L., and S.S. performed the connectomic analyses. A.M., S.S., and K.Y. interpreted the data and wrote the paper.

## Competing interests

The authors declare no competing interests.

## Data and materials availability

Correspondence and requests for materials should be addressed to S.S for connectome data or K.Y. for physiological data.

## Supplementary Materials

### Materials and Methods

#### Animals

Wild-type mice (C57BL/6J) were obtained from Janvier Labs. ChAT-IRES-Cre (strain: *Chat^tm2(cre)Lowl^*/MwarJ, Jackson laboratory stock: 028861) and Rosa-STOP-tdTomato (*Gt(ROSA)26Sor^tm9(CAG^-tdTomato)Hze*/J, Jackson laboratory stock: 007905) were used for bipolar cell imaging. Cart-IRES-Cre (strain: *Cartpt^tm1.1(cre)Hze^*/J, Jackson laboratory stock: 028533) was used for DSGC dendrite imaging. Oxtr-T2A-Cre; Thy1-STOP-EYFP (strain: *Cg-Oxtr^tm1.1(cre)Hze^*/J, Jackson laboratory stock: 031303; strain: Cg-Tg(Thy1-EYFP)15Jrs/J, Jackson laboratory stock: 005630) was used for electrophysiological recordings. Rosa-iDTR (strain: *Gt(ROSA)26Sor^tm1(HBEGF)Awai^*/J, Jackson laboratory stock: 007900) crossed with ChAT-Cre was used for the genetic ablation of starburst cells. These mice were purchased from Jackson laboratory and maintained in a C57BL/6J background. We used 8- to 16-week-old mice of either sex. Mice were group-housed throughout and maintained in a 12-hour/12-hour light/dark cycle with *ad libitum* access to food and water. All animal experiments were performed according to standard ethical guidelines and were approved by the Danish National Animal Experiment Committee (Permission No. 2015−15−0201−00541 and 2020-15-0201-00452).

#### Retinal preparation

Retinas were isolated from the left eye of mice dark-adapted for 1 hour before experiments. The isolated retina was mounted on a small piece of filter paper (MF-membrane, Millipore), in which a 2 × 2 mm window had been cut, with the ganglion cell side up. During the procedure, the retina was illuminated by dim red light (KL 1600 LED, Olympus) filtered with a 650 ± 45 nm band-pass optical filter (ET650/45×, Chroma) and bathed in Ringer’s medium (in mM): 110 NaCl, 2.5 KCl, 1 CaCl_2_, 1.6 MgCl_2_, 10 D-glucose, 22 NaHCO_3_ bubbled with 5 % CO_2_, 95 % O_2_. The retina was kept at 35-36°C and continuously superfused with oxygenated Ringer’s medium during recordings.

#### Electrophysiology

Electrophysiological recordings were conducted with an Axon Multiclamp 700 B amplifier (Molecular Devices). Signals were acquired using customized software on LabVIEW (National Instruments) developed by Zoltan Raics (SENS Software) and digitized at 10 kHz. Borosilicate glass micropipettes pulled by a micropipette puller (P-97, Sutter Instrument) were used for recordings. The firing discharges were recorded in cell-attached mode using pipettes filled with Ringer’s medium. To visualize the dendrites of recorded neurons, Alexa 594 was added in intracellular solution (in mM): 112.5 CsCH_3_SO_3_, 1 MgSO_4_, 7.8 × 10^−^3 CaCl_2_, 0.5 BAPTA, 10 HEPES, 4 ATP-Na_2_, 0.5 GTP-Na_3_, 5 QX314-Br, 7.5 neurobiotin chloride. pH was adjusted to 7.2 with CsOH. The equilibrium potential for chloride was calculated to be ~ −60 mV. The resistance of pipettes was 5-8 mOhm for cell-attached recording. The labeled cells were targeted using a two-photon microscope equipped with a mode-locked Ti: sapphire laser (Mai Tai DeepSee, Spectra-Physics), set to 940 nm, integrated into the physiological recording setup (SliceScope, Scientifica), as described previously (*18*). The two-photon fluorescence image was overlaid on the infra-red (IR) image acquired by a CCD camera (RT3, SPOT Imaging). The IR light was generated by a digital light projector (NP-V311X, NEC) with a 750 ± 25 nm filter.

For pharmacological experiments, we used SR95531 (50 μM, Sigma) to bock GABA_A_ receptors, TPMPA (100 μM, Sigma) to block GABA_C_ receptors, alpha-Bungarotoxin (0.1 μM, Tocris) to block α7-nACh receptors, and tetrodotoxin (1 μM, Tocris) to block Na^+^ channels. These agents were bath-applied during recordings.

#### AAV production

The production plasmid for AAV-8BP/2-shortCAG.SF-iGluSnFR.A184S-WPRE-SV40p(A) for bipolar cell imaging was developed by Zurich Viral Vector Core by transferring the insert from pAAV.CAG.SF-iGluSnFR.A184S (Addgene #106198). The plasmid encoding the AAV capsid 2/8BP2 (*14*) was kindly provided by Jean Bennett (University of Pennsylvania). The AAV was produced by Zurich Viral Vector Core (v454; 3.5 × 10^12^ GC/ml). AAV9.hSyn.Flex.iGluSnFr.WPRE.SV40 for ganglion cell imaging was obtained from Penn Vector Core (#98931; 7.73 × 10^13^ GC/ml).

#### Virus injections

Mice were anesthetized with an i.p. injection of fentanyl (0.05 mg/kg body weight; Actavi), midazolam (5.0 mg/kg body weight; Dormicum, Roche), and medetomidine (0.5 mg/kg body weight; Domitor, Orion) mixture dissolved in saline. We made a small hole at the border between the sclera and the cornea with a 30-gauge needle. The AAV was delivered through a pulled borosilicate glass micropipette (30 μm tip diameter). All pressure injections were performed using a Picospritzer III (Parker) under a stereomicroscope (SZ61; Olympus). For targeting bipolar cells, 1 μl was pressure-injected through the hole into the subretinal space of the left eye. For targeting ganglion cells, 2 μl was pressure-injected into the vitreous of the left eye. Mice were returned to their home cage after anesthesia was antagonized by an i.p. injection of flumazenil (0.5 mg/kg body weight; Anexate, Roche) and atipamezole (2.5 mg/kg body weight; Antisedan, Orion Pharma) mixture dissolved in saline and, after recovering, were placed on a heating pad for one hour.

#### Diphtheria toxin injections

To genetically ablate starburst amacrine cells, we used Rosa-iDTR (+/−) × ChAT-IRES-Cre mice and control Rosa-iDTR (-/-) × ChAT-IRES-Cre mice (fig. S4, A to D). Diphtheria toxin stock solution was made from diphtheria toxin (D0564, Sigma), which was dissolved in PBS (1 μg/μl), and stored at −80°C. Immediately before the injection, the injection solution (1 ng/μl) was prepared by diluting the stock solution in PBS. First, we subretinally injected AAV-8BP/2-shortCAG.SF-iGluSnFR.A184S-WPRE-SV40p(A) to target T7 and T2 bipolar cells for glutamate imaging. Nine days later, 2 μl diphtheria toxin was intravitreally injected into each of both eyes. The eyes were re-injected with the same amount of diphtheria toxin two days after the initial injection.

#### Two-photon glutamate imaging

Three to four weeks after virus injection, we performed two-photon glutamate imaging (*18*). The isolated retina was placed under the microscope (SliceScope, Scientifica) equipped with a galvo-galvo scanning mirror system, a mode-locked Ti: Sapphire laser tuned to 940 nm (MaiTai DeepSee, Spectra-Physics), and an Olympus 60× (1.0 NA) or Olympus 25x (1.05 NA) objective. The retina was superfused with oxygenated Ringer’s medium. The iGluSnFR signals emitted were passed through a set of optical filters (ET525/50m, Chroma; lp GG495, Schott) and collected with a GaAsP detector. Images were acquired at 8-12 Hz using custom software developed by Zoltan Raics (SENS Software). Temporal information about scan timings was recorded by TTL signals generated at the end of each scan, and the scan timing and visual stimulus timing were subsequently aligned during off-line analysis.

#### Visual stimulation

The visual stimulation was generated via custom-made software (Python and LabVIEW) developed by Zoltan Raics (SENS Software). For electrophysiological recordings, the stimulus was projected through a DLP projector (NP-V311X, NEC). The stimulus was focused on the photoreceptor layer of the mounted retina through a condenser (WI-DICD, Olympus). The intensity was measured using a photodiode power meter (Thorlabs), and the power of the spectrum was measured using a spectrometer (Ocean Optics). The calculated photoisomerization rate ranged from 0.0025 to 0.01 × 10^7^ photons absorbed per rod per second (R*/s) both for electrophysiological recordings and two-photon imaging. For glutamate imaging, the stimulus was projected using a DLP projector (LightCrafter Fiber E4500 MKII, EKB Technologies) coupled via a liquid light guide to an LED source (4-Wavelength High-Power LED Source, Thorlabs) with a 400 nm LED (LZ4-00UA00, LED Engin) through a band-pass optical filter (ET405/40×, Chroma). The stimuli were exclusively presented during the fly-back period of the horizontal scanning mirror (*18*). The contrast of visual stimulus (*C*_*S*_) was calculate as,

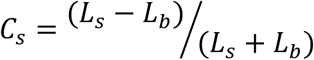

in which *L*_*S*_ and *L*_*b*_ indicate luminance intensity in stimulus and background, respectively.

#### Response measures

To evaluate sensitivity to luminance increments (ON) or decrements (OFF), we used static flash spots (300 μm in diameter, 2 s in duration, 100% positive contrast). To evaluate release kinetics, we used modulating spot (*2,18*). The stimulus (300 μm in diameter) had four phases: static flashing spot of 100% contrast, one of 50% contrast, one with increasing temporal frequency from 0.5 to 8 Hz, and one with increasing contrast from 5 to 80%.

To measure directional tuning and motion speed preference, we used a spot (300 μm in diameter, 100% positive contrast) moving in eight directions (0-315°, Δ45°) at 300 and 800 μm/s. The preferred direction was defined by a direction which elicited the maximum response, and the null direction was the opposite. To quantify the directional selectivity, we used a direction selectivity index (DSI) as,

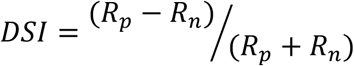

in which in which *R*_*p*_ and *R*_*n*_ indicate peak response amplitude in the preferred and null direction, respectively. DSI ranged from 0 to 1, with 0 indicating a perfectly symmetrical response, and 1 indicating a response only in the preferred direction.

The skewness (*Sk_w_*) in DSI histogram was calculated to quantify the potential biases in the distribution by mean-variance around the mode:

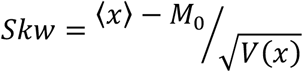

where *x* is DSI of individual input.*M*_0_ indicates a mode of distribution. < > and *V* () indicate mean and variance, respectively. To quantify precisely the directional biases in motion responses, an angle of preferred direction (*θ*) was defined by the vector sum:

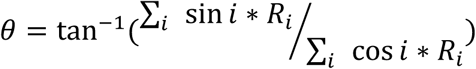

where *i* denotes the motion direction, and *R*_*i*_ denotes the response amplitude. For the analysis, we used ON-OFF DS ganglion cells whose DSIs were higher than 0.3 (9 of 12 cells: fig. S6D).

To quantify the effects of pharmacological blockers in light-evoked responses, we calculated change index (*CI*):

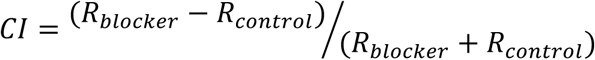

where *R_control_ R_blocker_* response amplitudes in control and after blocker application, respectively.

#### Regions of interests (ROIs) detection

ROIs for glutamate signals were determined by customized programs in MATLAB. First, the stack of acquired images was filtered with a Gaussian filter (3 × 3 pixels), and then each image was downsampled to 0.8 of the original using a MATLAB imresize function. The signals in each pixel were resampled using the MATLAB interp function with a rate of 2 and smoothed temporally by a moving average filter with a window size of 2 time-bin. Next, we calculated the temporal correlation among the pixels during static flash stimulus based on a raw cross-correlation 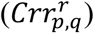:

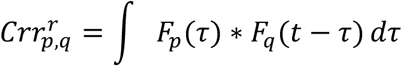

in which *F*_*p*_ and *F*_*q*_ indicate glutamate signals in pixel *p* and *q*, respectively. The noise correlation (*nc_p,q_*) was then given by a subtraction:

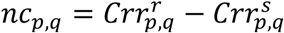

in which *CRR^S^* indicates trials-shuffled cross-correlation. The noise correlation was normalized as (*N C_p,q_*):

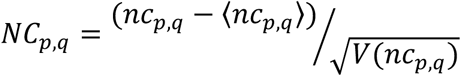

in which, < > and V () indicate mean and variance, respectively. We set a threshold of the score at 0 time-lag as 0.5 to determine which pixels were to be included as a single ROI. Then the response of each ROI (Δ*F*(*t*)) was calculated as

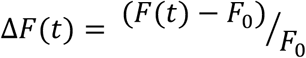

where F(*t*) is the fluorescent signal in arbitrary units, *F*_0_ is the baseline fluorescence measured as the average fluorescence in a 1-second window before the presentation of the stimulus. After the processing, responsive pixels were detected based on a response index (*RI*):

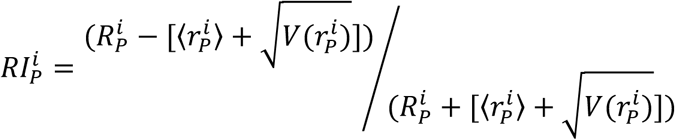

where *R^i^* is a peak response amplitude during motion stimulus to direction *i* in ROI *P*, and 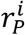 indicates glutamate signals before the stimulus (1 s period). The ROIs with the RI higher than 0.6 were determined as responsive.

#### Clustering

The clustering for classifying glutamate inputs to DS ganglion cells was based on the temporal kinetics in the responses to modulating spot, as described previously (*2,18*). We used a sparse principal component analysis (sPCA) to extract temporal features in response to a modulating flash based on the SpaSM toolbox on MATLAB (*28*). Next, we fitted a Gaussian mixture model based on the expectation-maximization algorithm using the MATLAB gmdistribution function to the dataset of detected sparse features. To determine the optimal number of clusters in the model, we calculated the Bayesian information criterion (*BIC*) score (*29*):

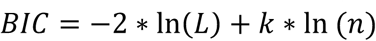

in which *L* is the log-likelihood of the model, *k* is the number of dimensions in the model, and *n* is the number of datasets. To separate ON and OFF input groups, we first performed the clustering using responses to static flash (the first phase in modulating spot), then we repeated the clustering, for the dissected each ON and OFF group, using the responses to the whole stimulus phases in the modulating spot. We sorted the detected clusters based on the similarity calculated by hierarchical clustering analysis using a standard linkage algorithm by MATLAB linkage function (Fig. 4E; fig. S6, G and I). We identified 12 groups (G1-G12; 6 ON and 6 OFF groups) in the glutamate inputs to ON-OFF DS ganglion cells.

#### Connectomic reconstruction

To characterize the bipolar-cell inputs to a DS ganglion cell and the amacrine-cell input to bipolar cells, we analyzed a set of serial electron microscopic sections of the adult mouse retina. The volume (k0725) is described in detail elsewhere (*24*). The volume was obtained from a young adult mouse (C57BL/6; 30 days of age) and fixed for 2 hr at room temperature in 2% buffered glutaraldehyde. A 1 mm^2^ sample obtained roughly midway between the optic disk and retinal margin was excised, stained with heavy metals to reveal synaptic ribbons and vesicles and other intracellular detail, dehydrated, and embedded in Epon Hard. A trimmed block (~200 mm x 400 mm) was imaged in a scanning electron microscope with a field-emission cathode (QuantaFEG 200, FEI Company). Back-scattered electrons were detected using a custom-designed detector and custom-built current amplifier. The incident electron beam delivered about 10 electrons/nm^2^. Imaging was performed at a high vacuum. Sides of the block were evaporation-coated with gold. The block face was serially cut as described elsewhere. Using a 26 nm section thickness, 10112 consecutive block faces were imaged, yielding aligned data volumes of 4992 x 16000 x 10112 voxels (135 mosaics of 3584 x 3094 images). This corresponds to a spatial volume of approximately 66 x 211 x 263 mm. The smallest dimension corresponds to retinal depth, which ranged from the ganglion cell layer to the innermost part of the inner nuclear layer. The edges of neighboring mosaic images overlapped by ~1 mm. Mosaics and slices were aligned offline to subpixel precision by Fourier shift-based interpolation. The datasets were then split into cubes (128 x 128 x 128 voxels) for import into Knossos (http://knossostool.org), a freely available software package for exploration and skeletonization of cell profiles in SBEM datasets. We also used webKnossos (http://webknossos.org), a cloud- and browser-based 3D annotation tool for large-scale data analysis of SBEM data.

#### Histology and confocal imaging

After the two-photon imaging experiments, retinas were fixed for 30 minutes in 4% paraformaldehyde in PBS and washed with PBS overnight at 4 ? on a shaker. The retinas were incubated in 30% sucrose in PBS for at least 3 hours at room temperature (RT). To enhance the penetration of antibodies, retinas were transferred in the sucrose buffer and frozen and thawed three times. After washing with PBS, retinas were blocked for 3 hours in blocking buffer (1% bovine serum albumin [BSA], 10% normal donkey serum [NDS], 0.5% TritonX 100, 0.02% sodium azide in PBS) at RT. The retinas were incubated with primary antibodies (chicken anti-GFP 1:1000 [abcam, ab13970]; goat anti-ChAT 1:200 [Milipore, ABN1144P]) for 5 days at RT in antibody reaction buffer (1% BSA, 3% NDS, 0.5% TritonX 100, 0.02% sodium azide in PBS), and secondary antibodies (donkey anti-chicken IgY Alexa 488 1:200 [Jackson ImmunoResearch, AB 2340375]; donkey anti-goat IgG Alexa 568 1:200 [Invitrogen, A11057]) for one day at 4 ? in antibody reaction buffer. After a final washing in PBS, retinas were embedded in Fluoromount-G (eBioscience). The stained retinas were imaged using a confocal microscope (Zeiss LSM 780) using a 40x (1.4 NA) or 63x (1.2 NA) objectives. The images were acquired at 1024 × 1024 pixel (0.35 μm/pixel for 40x; 0.22 μm/pixels for 63x), and the optical thickness of each imaging plane in z-stack was 0.3 μm. The images were processed and analyzed using MATLAB.

#### Statistical analysis

All analyses and statistical tests were performed by MATLAB 2017b (Mathworks). Population data were shown as mean ± SD. To compare the differences in paired conditions, Wilcoxon singed-rank test was used. To compare the differences in different groups, Mann-Whitney-Wilcoxon test was used. To compare the differences in distribution, Kolmogorov-Smirnov test was used. To evaluate the uniformity in angler distribution, Hodges-Ajne test was used. Fitting of the Gaussian function was based on the least-square method in MATLAB (Fig. 4; fig. S2). The numbers of modality in the preferred direction of bipolar cell axonal boutons (Fig. 1; fig. S2) were evaluated by Silverman’s test using kernel density estimation. The kernel was a Gaussian kernel. The p-values were calculated by the Bootstrap method (2000 times replication; fig. S2).

#### Data and code availability

The datasets and code generated in this research are available from the corresponding author, Keisuke Yonehara for imaging data or Shai Sabbah for EM data upon reasonable requests.

---

**Fig. S1.**
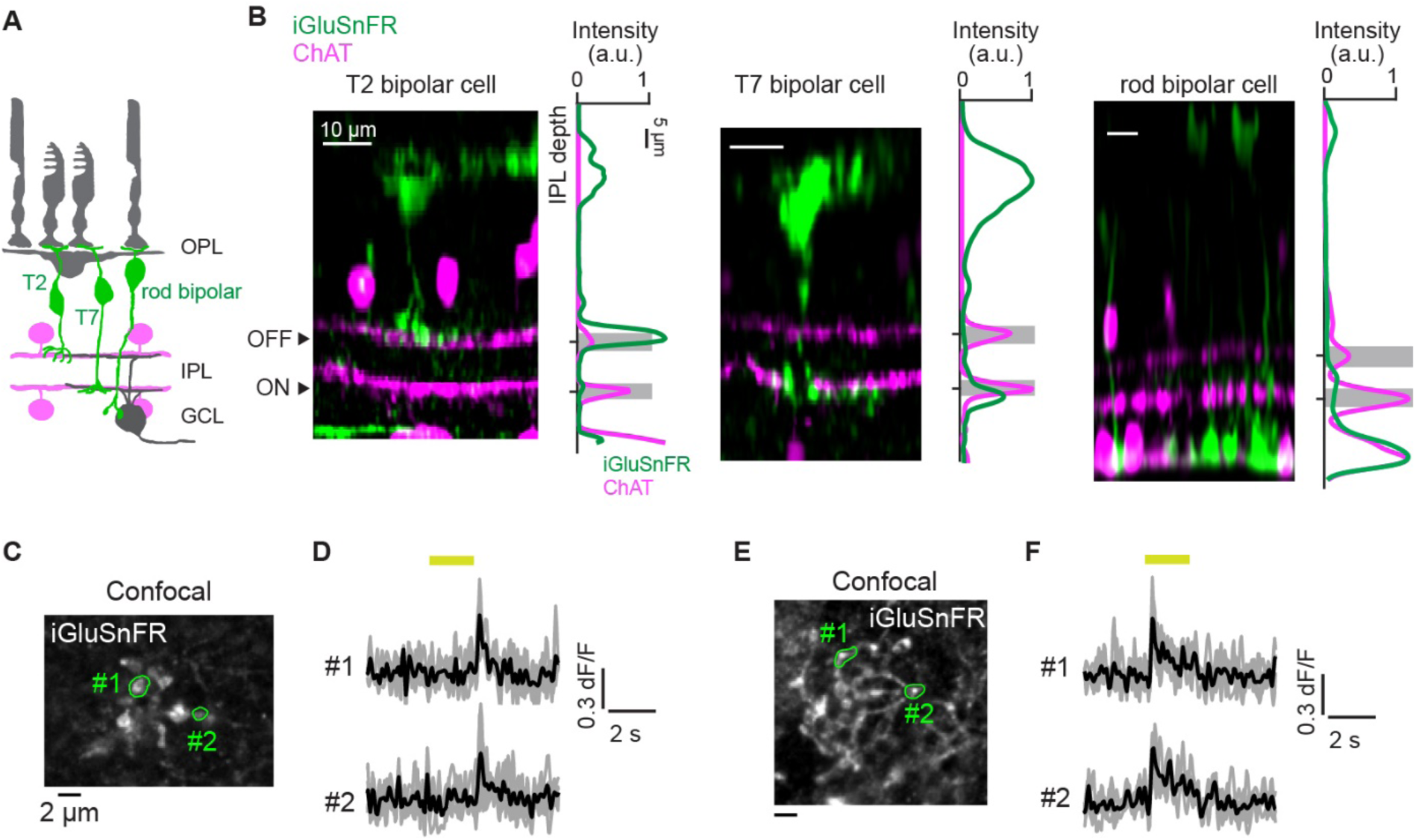
Morphology of the bipolar cells labeled by SF-iGluSnFR.A184S. (**A**) A schematic for labeled bipolar cells (green; type 2 [T2], type 7 [T7], and rod bipolar cells) and starburst cells (magenta). OPL, outer plexiform layer. IPL, inner plexiform layer. GCL, ganglion cell layer. (**B**) Example cross-sectional views of confocal reconstructions of the labeled bipolar cells and the cross-sectional intensity of the fluorescence of iGluSnFR (green) and ChAT (magenta). Gray bands, depth of ON and OFF ChAT signals. We found one OFF bipolar cell type whose axon terminals were stratified slightly above the OFF ChAT band (left) and one ON bipolar cell type whose axon terminals were stratified slightly below the ON ChAT band (center), supporting the morphological features of type 2 and 7 bipolar cells, respectively (*1, 3, 16*). Another labeled bipolar cell type had large axon terminals at the layer of starburst cell’s soma (right), suggesting it is rod bipolar cell. (**C and D**) Confocal image of axonal boutons of type 2 bipolar cells (C) and the light-evoked responses obtained by the two-photon imaging in the corresponding field-of-view (D). Black and gray, an average and each trial, respectively. Yellow line, the timing of light stimulus (2 s static flash). (**E and F**) Confocal image of axonal boutons of type 7 bipolar cells (E) and the light-evoked responses (F).

**Fig. S2.**
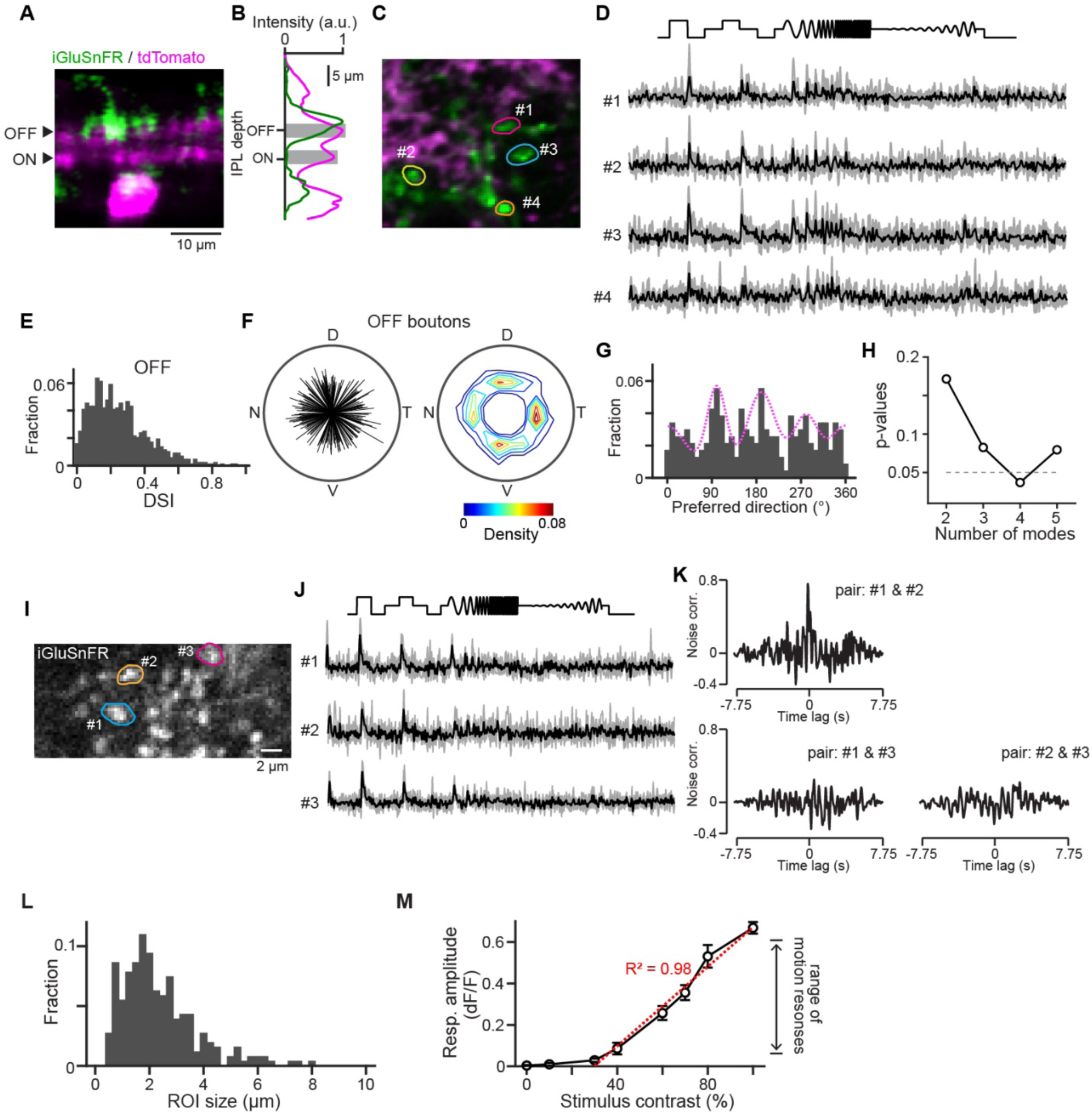
Two-photon imaging from the labeled ON and OFF bipolar cell axon terminals. (**A**) Example two-photon image showing the side view of SF-iGluSnFR.A184S-labeled bipolar cells (green) and tdTomato-labeled starburst processes (magenta) focusing on the area in which OFF bipolar terminal is evident. (**B**) Intensity of iGluSnFR (green) and tdTomato (magenta) fluorescence (A) along the depth. Gray bands, depth of ON and OFF tdTomato signals. (**C**) Example tangential view of iGluSnFR signals in the OFF layer. (**D**) Light-evoked responses in regions of interests (ROIs) shown in c. Gray, each trial. Black, average. (**E**) Histograms of DSI in OFF boutons. Mean ± SD, 0.25 ± 0.17, 881 OFF boutons. (**F**) Polar plots of directional tunings in individual OFF boutons (left; length, DSI; angle, preferred direction) and the density plots (right). (**G**) Histogram of the preferred direction. Magenta curve, a fitted Gaussian function. (**H**) Summary of the calculated *p*-values in Silverman’s test for multimodality in the preferred directions of boutons. (**I and J**) Example FOV of ON layer (I) and responses to modulating spot in three axonal boutons (J; #1-#3, colored circles in I). Black and gray, responses in an average and four trials, respectively. All three boutons have similar response profiles, indicating that the three belong to the same functional group. (**K**) Noise correlation during static flash (Materials and Methods) in the pairs among the three boutons. A pair of #1 and #2 showed high noise correlation at 0 time-lag, indicating the two boutons belong to the same bipolar cell. Boutons with noise correlation higher than 0.5 were assigned to the same bouton cluster (see Fig. 1). (**L**) Histogram of ROI size. (**M**) Relationship between stimulus intensity and response amplitudes. The range of motion response amplitudes that were used in this study showed strong linearity (R^2^ = 0.98 in a linear fitting).

**Fig. S3.**
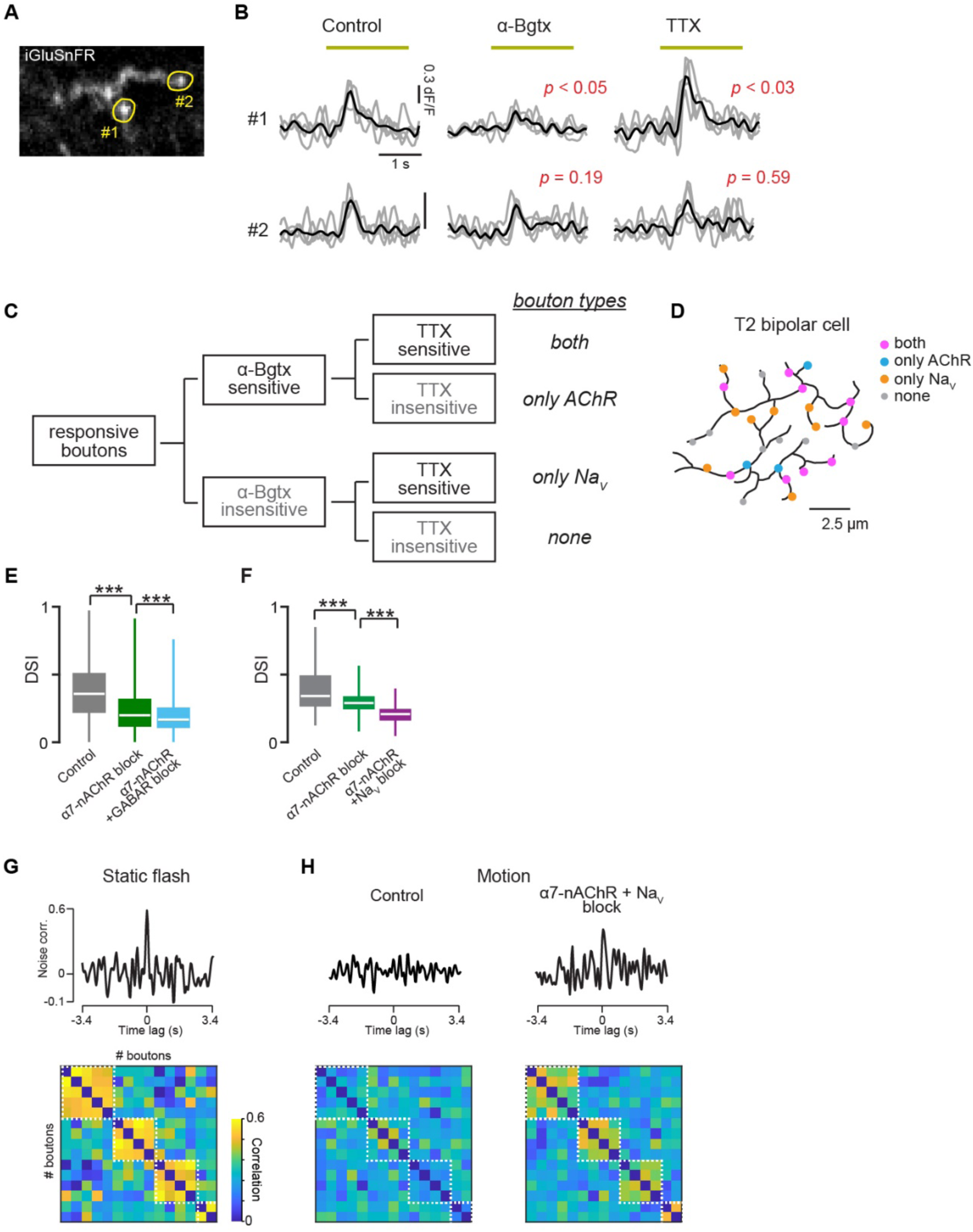
Different modulation of bipolar cell axonal boutons. (**A and B**) Example FOV of ON layer showing axonal boutons (A; #1 and #2) and their light-evoked responses to static flash (yellow line, 300 μm diameter, 2 s duration, 100% contrast) in control, α7-nAChR blocking (α-Bgtx), and Na_V_ blocking (TTX) conditions. Gray and black, responses in four trials and an average, respectively. The *p*-values indicate the results of a two-tailed *t*-test for peak response amplitudes in each trial between control and blocker conditions. (**C**) Schematic for the classification of boutons based on the differences of sensitivity to α7-nAChR and Na_V_ blockings based on the statistical significance in the changes of response amplitudes by applications of the blockers (B). (**D**) Bouton types on a T2 bipolar cell axon. (**E**) Changes of DSI in ON and OFF DS boutons by α7-nAChR blocking (green) and additional GABA receptor blockings (cyan; SR+TPMPA). 82 boutons. (**F**) Changes of DSI in ON and OFF DS boutons by α7-nAChR blocking (green) and additional Na_V_ blockings (purple; TTX). 242 boutons. (**G**) Top, noise correlation in an example pair of boutons belonging to the same bipolar cell during a static flash. Bottom, correlation matrix showing noise correlation (correlation at 0 time-lag) in a static flash. White dotted squares indicate bouton groups sharing high noise correlation in a static flash. (**H**) Noise correlation (top) and the correlation matrix (bottom) during motion stimulus in control (left) and after blocking of α7-nAChR and Na_V_ (right). The correlated bouton activities were decorrelated in motion stimulus, and the decorrelation was impaired by the blockings of α7-nAChR and Na_V_. ***, *p* < 0.001.

**Fig. S4.**
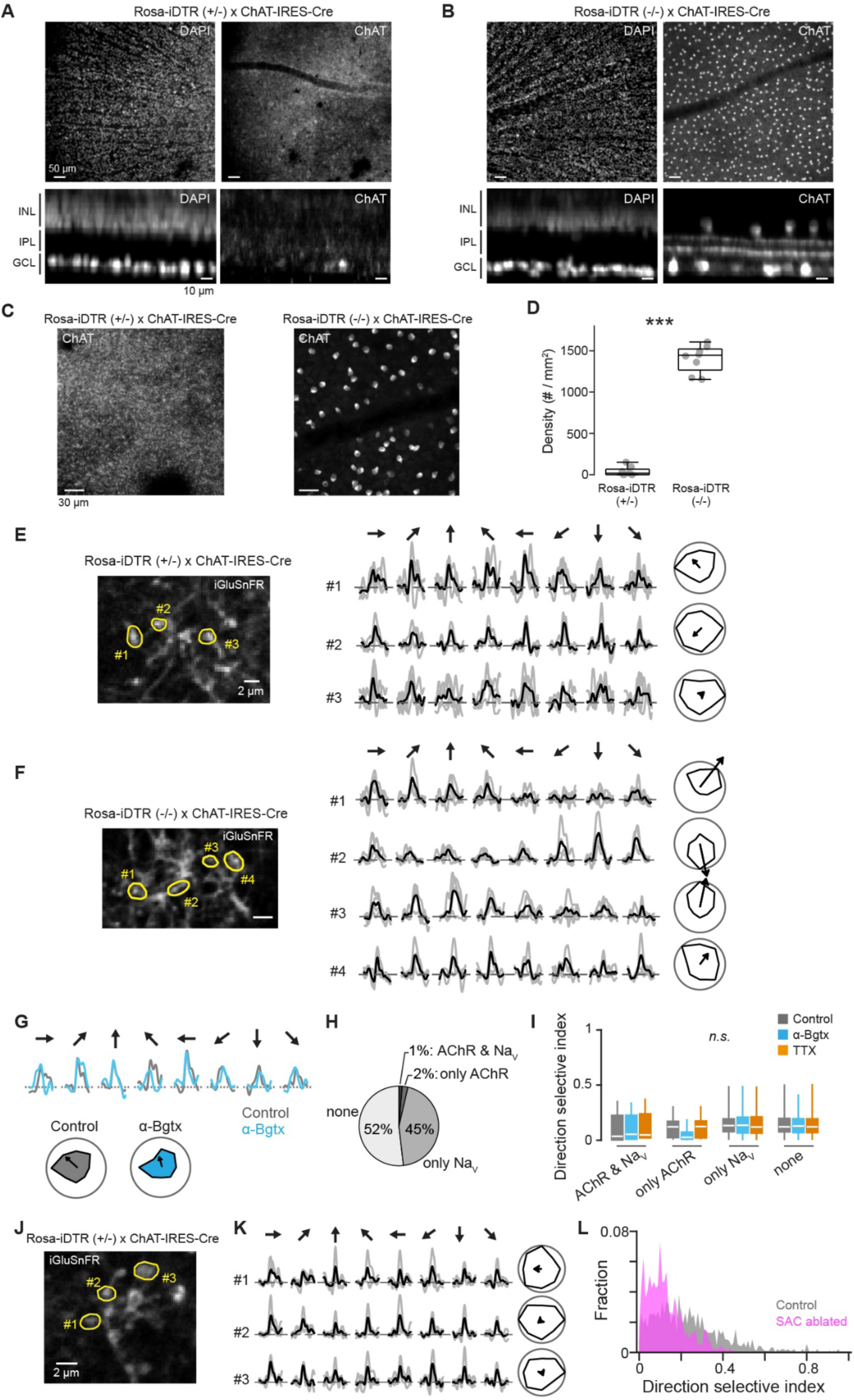
Ablation of starburst cells impaired direction selectivity in T7 and T2 bipolar cells. (**A and B**) Top, example confocal images of DAPI (left) and ChAT (right) in Rosa-iDTR (+/−) x ChAT-IRES-Cre (A) and Rosa-iDTR (-/-) × ChAT-IRES-Cre (B) retina after injection of diphtheria toxin. Bottom, the cross-sectional views of confocal reconstructions. INL, inner nuclear layer. IPL, inner plexiform layer. GCL, ganglion cell layer. ChAT in INL of Rosa-iDTR (+/−) retina did not show clear signals. (**C and D**) ChAT signals in GCL in Rosa-iDTR (+/−) (left) and Rosa-iDTR (-/-) (right) retina (C) after injection of diphtheria toxin, and the comparison of calculated starburst cells density. Gray circles, 16 images from 5 Rosa-iDTR (+/−) and 4 Rosa-iDTR (-/-) retina injected with diphtheria toxin. (**E and F**) Example FOVs of ON layer showing axonal boutons (left) and their light-evoked responses to motion stimulus (right) in Rosa-iDTR (+/−) (E) and Rosa-iDTR (-/-) (F) retina. Gray and black, responses in four trials and an average, respectively. (**G**) Example responses to motion stimulus (top) and the directional tunings (bottom) in control (gray) and blockades of α7-nAChR (cyan) of a bouton in Rosa-iDTR (+/−) (#1 in E). (**H**) The population of bouton types with distinct sensitivity to the blockades of α7-nAChR and Na_V_ in Rosa-iDTR (+/−). (**I**) Summary of DSI changes by the blockades of α7-nAChR (cyan) and Na_V_ (orange) in Rosa-iDTR (+/−). 5 both, 9 only AChR, 172 only Na_V_, 200 none boutons. (**J and K**) Example FOVs of OFF layer showing axonal boutons (J) and their light-evoked responses to motion stimulus (K) in Rosa-iDTR (+/−) retina. Gray and black, responses in four trials and an average, respectively. (**L**) Histograms of DSI in T2 terminals in Rosa-iDTR (-/-) (gray; 506 boutons, 4 retina) and Rosa-iDTR (+/−) (magenta; 473 boutons, 5 retina) retina. ***, *p* < 0.001. Mann-Whitney-Wilcoxon test.

**Fig. S5.**
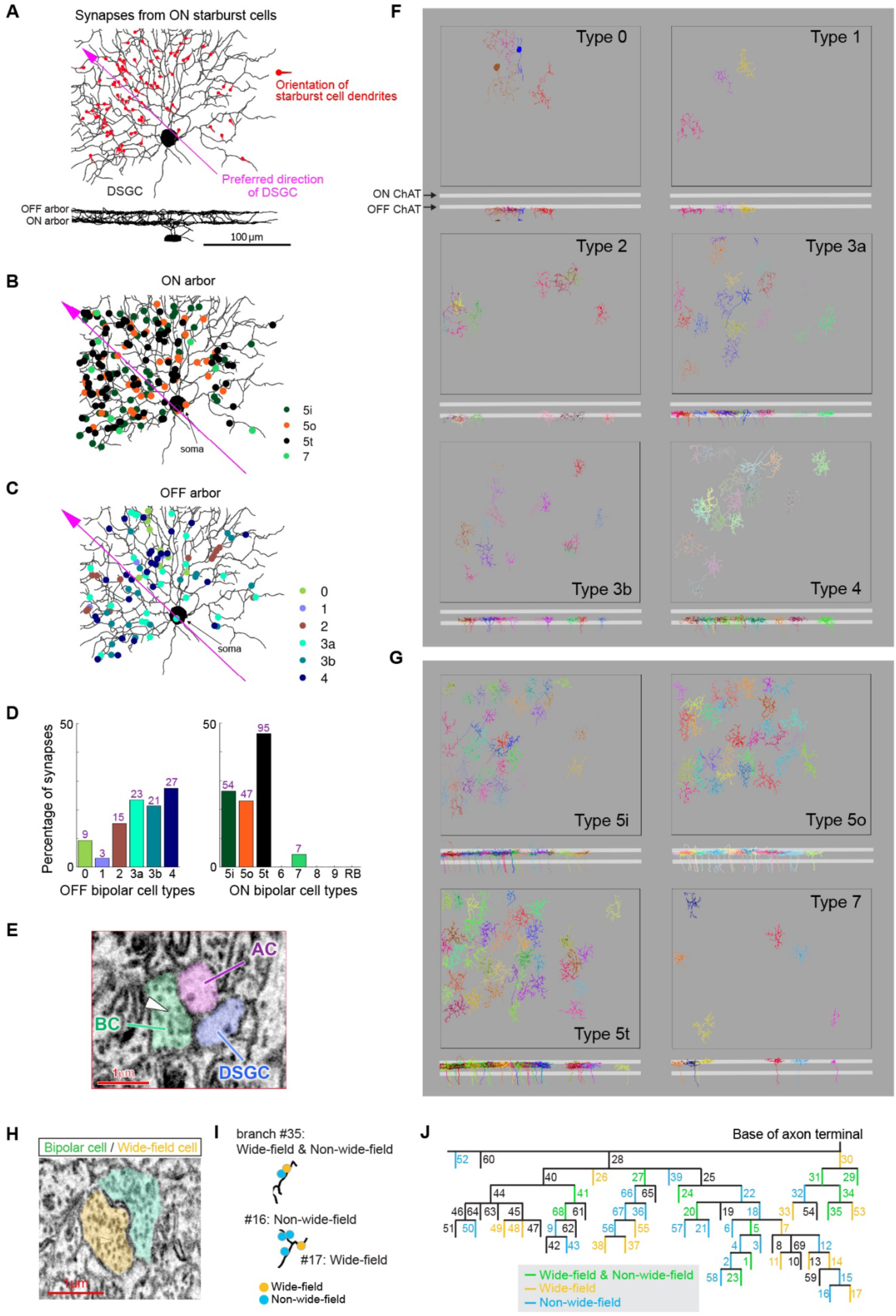
Bipolar cell inputs to ON-OFF DSGCs. (**A**) An SBEM reconstruction of an ON-OFF DSGC, at *en face* (top) and orthogonal (bottom) views. ON starburst cell wiring with the DSGC is asymmetric. Synapses between 57 ON starburst cells and the DSGC are marked (red pointers), with each pointer indicating the centrifugal orientation of the presynaptic starburst cell dendrite (i.e., away from the soma). The pointer’s tip marks the location of the synapse. The preferred direction vector (magenta) marks the average orientation opposite to the orientation of presynaptic starburst cell dendrites. The mapping of synapses and bipolar cell type identification was generously given and assisted by David Berson (Brown University). (**B and C**) Spatial distribution of bipolar cell-to-DSGC synapses, for the ON (B) and OFF (C) DSGC arbor. Synapses (filled circles) are color-coded based on the bipolar cell type forming the synapse. The preferred direction vector is marked by the arrow (magenta). (**D**) Percentage of ON (right) and OFF (left) bipolar cell inputs provided by each type. RB, rod bipolar cell. (**E**) An SBEM micrograph illustrating a dyad ribbon synapse between a bipolar cell axon terminal (BC, green), a postsynaptic amacrine cell (AC, purple), and a postsynaptic dendrite of the DSGC (blue). Arrowhead marks the synaptic ribbon that appears brighter than synaptic vesicles. (**F and G**) The different OFF (F) and ON (G) bipolar cell types in the plane of the retina (top), and at the orthogonal view (bottom), relative to the ON and OFF ChAT bands (two horizontal white stripes). (**H**) Example input synapses made from wide-field amacrine cells (yellow) on terminal branches of T7 bipolar cell (light green). (**I**) Example T7 terminal branches receiving inputs either from both wide-field (yellow circle) and non-wide-field (cyan circle) cells (top; branch#35), wide-field (bottom; branch#16), or non-wide-field (bottom; branch#17) cell. (**J**) A hierarchical tree of terminal branches of a T7 bipolar cell (Fig. 3D). Branches that formed a synapse only from wide-field (yellow), only from non-wide-field (cyan), and both wide-field and non-wide-field (green). The numbers indicate the branch IDs in Figure 3D.

**Fig. S6.**
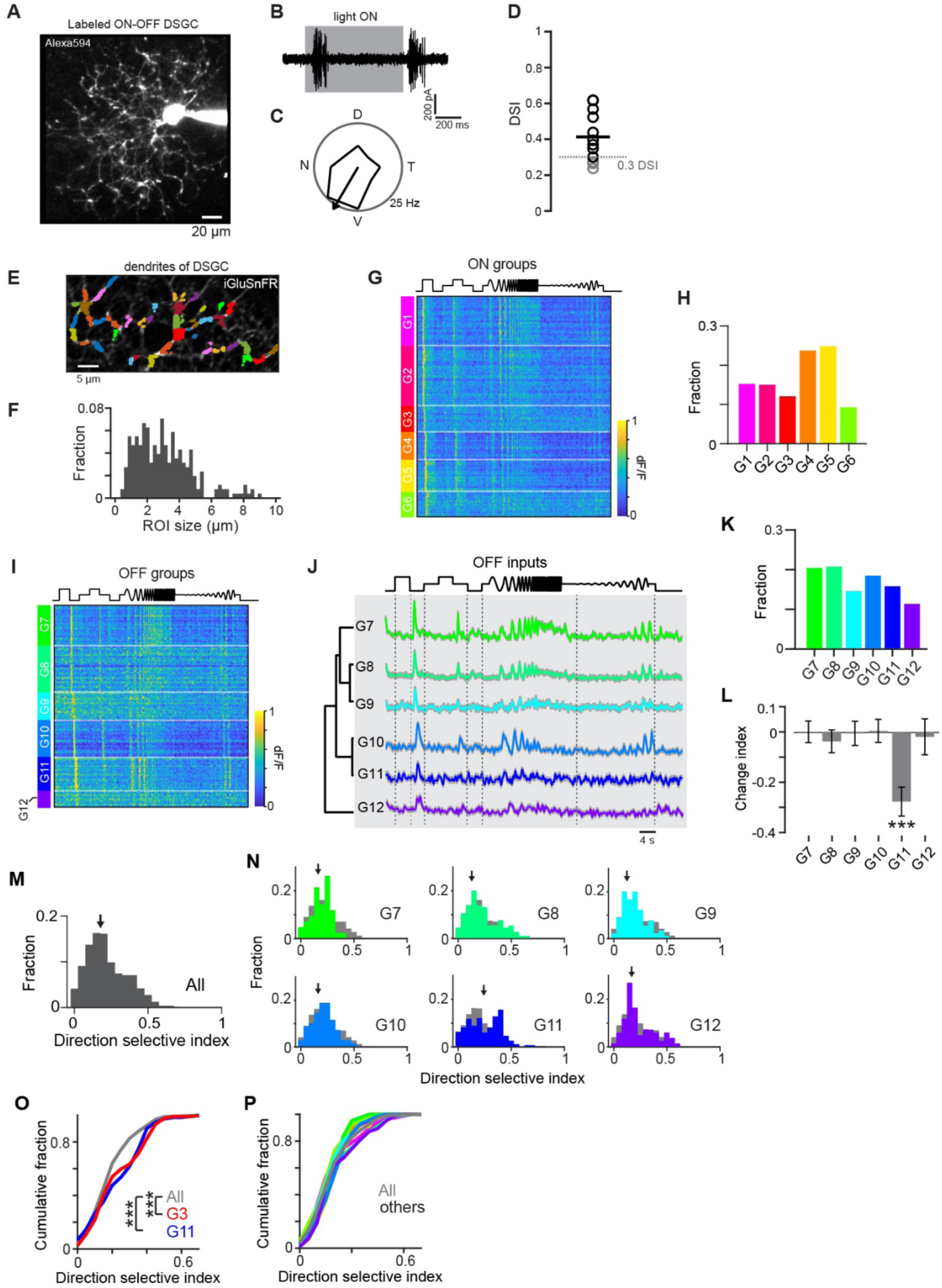
Characterization of glutamate inputs to ON-OFF DSGC dendrites. (**A**) A two-photon image of an ON-OFF DSGC filled with Alexa594 dye. The ON-OFF DSGCs were targeted by Cre-dependent iGluSnFR signals expressed in Cart-IRES-Cre mice. (**B**) Firing activity of the ON-OFF DSGC (A) to static flash as indicated by gray background (300 μm diameter, 2 s duration, 100% contrast). (**C**) Polar plot of firing activity of the ON-OFF DSGC (A) in response to a moving spot (C; 800 μm/s, 300 μm diameter, 100% contrast). An arrow indicates the preferred direction of this cell as determined by the vector sum. (**D**) DSI for recorded 12 cells (circles). We used 9 cells (black circles) with DSIs higher than 0.3 (gray dotted line) for subsequent glutamate imaging experiments. (**E**) An example FOV for two-photon glutamate imaging from ON dendrites of an ON-OFF DSGC. The defined ROIs were shown in different colors. (**F**) Histogram of ROI size for imaging of DSGC dendrites. (**G**) Responsive ROIs sorted by the identified ON input groups (G1-G6) based on the temporal features in responses to a modulating spot (*19*). The same clustering analyses were performed for OFF glutamate inputs as well (shown in I). (**H**) The fraction of ON groups. (**I to K**) Responsive ROIs sorted by the identified OFF input groups (I; G7-G12), average light-evoked responses in the six OFF groups (J), and their fraction (K). (**L**) Change index showing the effects of α7-nAChR blocking (α-Bgtx) in response amplitudes to static flash for OFF (G7-G12) group. 28 G7, 23 G8, 32 G9, 35 G10, 31 G11, 18 G12 inputs. (**M and N**) Summary of DSI histogram in all ON and OFF inputs (M) and OFF groups (N). Gray and colored, all inputs and individual groups, respectively. Arrows, median. (**O and P**) Cumulative histograms of DSI in G3 and G11 groups (O), and others (P). G3 (red) and G11 (blue) showed significantly different distribution from that of all inputs (gray). ***, *p* < 0.001. Wilcoxon singed-rank test for fig. S6L; Kolmogorov-Smirnov test for fig. S6, O and P.

**Fig. S7.**
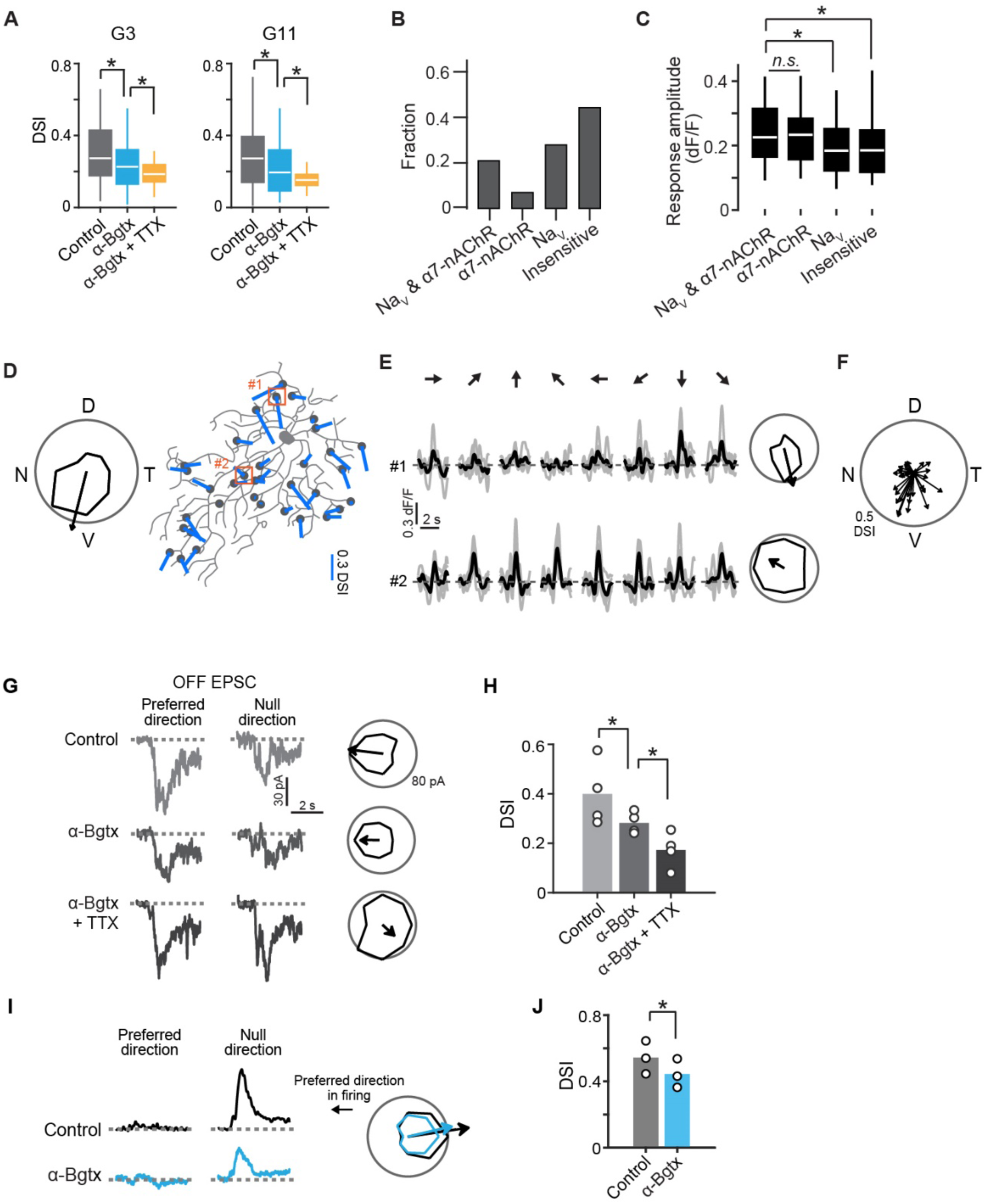
The effects of the tuned glutamate inputs to the excitatory and inhibitory inputs to ON-OFF DSGCs. (**A**) Effects of α7-nAChR and Na_V_ blocking on the directional tunings in G3 and G11 groups. 55 G3 and 72 G11, respectively. (**B**) The fraction of the four groups in G3 inputs: inputs sensitive to both Na_V_ and α7-nAChR blockings, only to α7-nAChR, only to Na_V_, and insensitive inputs. (**C**) Response amplitudes in the preferred direction motion for 32 Na_V_ and α7-nAChR-sensitive, 14 only α7-nAChR-sensitive, 48 only Na_V_-sensitive, and 76 insensitive inputs. The inputs with α7-nAChR sensitivity showed higher response amplitude than those in other input groups. (**D**) Left, directional tuning of firing activity in a ventral-direction-tuned ON-OFF DSGC. Right, the morphology of the ON-OFF DSGC and locations of glutamate inputs by G3 and G11 groups (black circles). Blur bars, vector sum in the directional tunings in individual inputs (length, direction selective index; angle, preferred direction). Orange squares, the locations of ROIs in E. (**E**) Tuned (#1) and untuned (#2) glutamate inputs to the ON-OFF DSGC. The tuned inputs were aligned to the firing preferred direction of the DSGC. (**F**) A polar plot for the directional tuning in the individual G3 and G11 groups on the ventral-direction-tuned DSGC (length, direction selective index; angle, preferred direction). (**G**) Excitatory postsynaptic currents (EPSC) evoked by the trailing edge of the spot (OFF motion stimulus) moving in the preferred and null direction of an ON-OFF DSGC in control, α-Bgtx, and TTX conditions. (**H**) DSI of EPSC induced by OFF motion stimulus in control, α-Bgtx, and TTX conditions. Bars, an average of 4 ON-OFF DSGCs (circles). (**I**) Inhibitory postsynaptic currents (IPSC) evoked by the spot moving in the preferred and null direction of ON-OFF DSGC’s firings. Black, control; cyan, α-Bgtx; orange, α-Bgtx, and TTX. (**J**) Effects of α7-nAChR blockings (α-Bgtx) on the directional tuning in IPSC. *, *p* < 0.05. Mann-Whitney-Wilcoxon test for fig. S7, A and C; Wilcoxon singed-rank test for fig. S7, H and J.

## Notes

### Competing Interest Statement

The authors have declared no competing interest.

## References and Notes

1. T. Euler, S. Haverkamp, T. Schubert, T. Baden, Nat Rev Neurosci. 15, 507–519 (2014).

2. K. Franke et al., Nature. 542, 439–444 (2017).

3. K. Shekhar et al., Cell. 166, 1308–1323.e30 (2016).

4. D. I. Vaney, B. Sivyer, W. R. Taylor, Nat Rev Neurosci. 13, 194–208 (2012).

5. S. Sabbah et al., Nature. 546, 492–497 (2017).

6. T. Euler, P. B. Detwiler, W. Denk, Nature. 418, 845–852 (2002).

7. K. L. Briggman, M. Helmstaedter, W. Denk, Nature. 471, 183–188 (2011).

8. K. Yonehara et al., Nature. 469, 407–410 (2011).

9. K. Yonehara et al., Neuron. 79, 1078–1085 (2013).

10. S. J. H. Park et al., J. Neurosci. 34, 3976–3981 (2014).

11. M. Chen et al., J. Neurophysiol. 112, 1950–1962 (2014).

12. K. A. Percival, S. Venkataramani, R. G. Smith, W. R. Taylor, J. Comp. Neurol. 527, 270–281 (2019).

13. S. Sethuramanujam et al., bioRxiv, (2020). doi:10.1101/2020.04.18.048330.

14. L. M. Hall, C. B. Hellmer, C. C. Koehler, T. Ichinose, Invest. Ophthalmol. Vis. Sci. 60, 1353–1361 (2019).

15. T. Cronin et al., EMBO Molecular Medicine. 6, 1175–1190 (2014).

16. Y. Tsukamoto, N. Omi, Front. Neuroanat. 11 (2017), doi:10.3389/fnana.2017.00092.

17. L. Liang et al., Cell. 173, 1343–1355.e24 (2018).

18. A. Matsumoto, K. L. Briggman, K. Yonehara, Current Biology. 29, 3277–3288.e5 (2019).

19. D. P. Lukasiewicz, Progress in Brain Research (Elsevier, 2005), vol. 147, pp. 205–218.

20. S. Lee, K. Kim, Z. J. Zhou, Neuron. 68, 1159–1172 (2010).

21. X. Huang, M. Rangel, K. L. Briggman, W. Wei, Nat Commun. 10, 1–15 (2019).

22. T. Baden, F. Esposti, A. Nikolaev, L. Lagnado, Current Biology. 21, 1859–1869 (2011).

23. S. Saszik, S. H. DeVries, J. Neurosci. 32, 297–307 (2012).

24. H. Ding et al, Nature. 535, 105–110 (2016).

25. S. Sabbah et al., bioRxive, (2018). doi:10.1101/442954.

26. E. S. Yamada et al., J. Comp. Neurol. 461, 76–90 (2003).

27. J. N. Kay et al, J. Neurosci. 31, 7753–7762 (2011).

28. H. Zou, T. Hastie, R. Tibshirani, J. Computational and Graphical Statistics. 15, 265–286 (2006).

29. C. Fraley, A. E. Raftery, J. the American Statistical Association. 97, 611–631 (2002).

30. J. S. Kim et al., Nature. 509, 331–336 (2014).

31. M. J. Greene, J. S. Kim, H. S. Seung, Cell Reports. 14, 1892–1900 (2016).

32. Z. Shi et al, J. Neurosci. 31, 13118–13127 (2011).

33. H. von Gersdorff, T. Sakaba, K. Berglund, M. Tachibana, Neuron. 21, 1177–1188 (1998).

